# Structural Basis for MCAM Mediated Inhibition of Laminin Dependent Cell Migration

**DOI:** 10.64898/2026.09.11.751044

**Authors:** Tongqing Li, Sofia Alarcon-Frias, Hengyi Li, Claudio R. Alarcón, Daryl E. Klein

## Abstract

MCAM/CD146 is an immunoglobulin-superfamily receptor expressed on endothelial cells, immune cells and numerous carcinomas. Although MCAM has long been associated with metastatic progression, recent work has revealed context-dependent inhibitory effects on transendothelial migration, leaving unresolved how MCAM engagement of basement membrane laminins alters cell migration. Here, we report multiple cryo-electron microscopy structures of human MCAM bound to laminin α4. We show MCAM engages laminin on a surface centered on the first two laminin globular domains (LG1-2) and that this epitope directly overlaps with the interface used by α6β1 integrin. Consistent with structural competition, MCAM occupation of laminin α4 abolishes integrin-dependent cell migration. Structure-guided mutation of the MCAM-laminin interface disrupts laminin binding and eliminates MCAM-mediated inhibition. These findings establish that MCAM and integrin compete for a shared interface on laminin α4 and define a structural mechanism by which MCAM occupancy can suppress laminin-dependent migration. The structure resolves a central ambiguity in MCAM biology and provides a framework for developing biologics that selectively modulate pathological cell migration.

## Introduction

Metastasis remains the principal cause of cancer-related mortality and requires tumor cells to traverse vascular and stromal barriers during dissemination[1, 2]. A central step in this process is transendothelial migration, during which tumor cells cross the endothelial monolayer and penetrate the underlying basement membrane[3]. Basement membranes regulate adhesion, traction, survival and migration, thereby influencing whether cells can efficiently breach tissue boundaries[4, 5].

Laminins are heterotrimeric glycoproteins that form a major architectural and signaling scaffold of basement membranes[4, 5]. Laminins are composed of three chains (α, β, γ) and named accordingly (i.e. Laminin-αβγ). Distinct laminin isoforms confer different adhesive and migratory properties. Laminin-411 and laminin-511 are major components of endothelial basement membranes, with laminin α4-containing matrices generally associated with more permissive immune-cell and tumor-cell migration, whereas laminin α5-containing matrices can impose more restrictive barrier properties[6, 7]. These functional differences are particularly relevant at post-capillary venules, which are major sites of leukocyte extravasation and tumor-cell vascular transit[6-8].

Integrins are the principal force-generating receptors that enable cells to adhere to, remodel and migrate across laminin-rich matrices[8, 9]. Laminin-binding integrins, including α6β1, engage laminin α globular domains and mechanically couple extracellular matrix engagement to the actin cytoskeleton[9, 10]. Recent structural work on α6β1 bound to laminin-511 established how the integrin headpiece recognizes a composite laminin surface involving the α-chain LG region and the γ-chain tail[10]. However, how other receptors engage α4-containing laminins, and whether those interactions are compatible with integrin binding, has remained unresolved.

MCAM, also known as CD146 or MUC18, is a type I transmembrane immunoglobulin-superfamily receptor expressed by endothelial cells, pericytes, activated lymphocytes and multiple tumor types[11-14]. It was originally identified as a melanoma cell adhesion molecule (MCAM) and has been widely linked to melanoma progression, vascular interaction and metastatic potential[11-14]. Consistent with this view, MCAM has been described as a receptor that promotes tumor-cell adhesion, migration and invasion in several settings[12-14]. In immune biology, MCAM marks inflammatory T-cell subsets and binds laminin-411, facilitating entry of TH17 cells into the central nervous system[15]. MCAM and soluble MCAM have also been implicated in monocyte transendothelial migration and angiogenic signaling[16, 17].

Yet MCAM biology has remained controversial. In breast cancer, MCAM has been reported to suppress tumor-cell adhesion to endothelial cells and inhibit transendothelial migration[18]. Mannion and colleagues further emphasized that MCAM expression in tumors is heterogeneous and may reflect malignant cells, vasculature, normal epithelium or EMT-associated cell states, complicating interpretation of bulk expression data[18]. Thus, the field has lacked a molecular framework to explain how MCAM can be associated with pro-migratory phenotypes in some contexts while inhibiting migration in others. A critical unresolved question is whether MCAM and integrins cooperate on laminin substrates or instead compete for the same laminin surface. Previous biochemical studies showed that α6β1 integrin and MCAM bind α4-containing laminins and suggested that their binding sites lie near one another on the laminin α4 globular domain[19, 20]. However, no high-resolution structure has visualized MCAM bound to laminin α4, and no structural mechanism has explained whether MCAM binding is compatible with integrin engagement.

Here we define the MCAM-laminin α4 interaction by cryo-EM and show that MCAM directly occludes the α6β1 integrin-binding surface. Functional and mutational analyses demonstrate that this structural interface is required for MCAM-mediated inhibition of laminin-dependent migration. These findings establish a direct competitive mechanism for MCAM and provide a unifying framework for interpreting context-dependent MCAM function in migration and metastasis.

## Results

### Cryo-EM structure of the MCAM-Laminin α4 complex

To study the interaction between MCAM and LamA4, we purified multiple recombinant constructs of both proteins (**Supplementary Fig. 1a**). For LamA4 globular domain expression (LG1-3) we used either a construct that maintained a minimal heterotrimer coiledcoil designated T8[21], or the alpha chain’s globular domains in isolation without the beta or gamma chains (Fig. 1a). For MCAM expression we used a soluble construct (sMCAM, residues G26–G559) that comprises all five extracellular immunoglobulin (Ig)-like domains: two N-terminal V-type domains (V1, V2) followed by three C2-type domains (C1, C2 and C3) (**Fig. 1a**).

**Figure 1.**
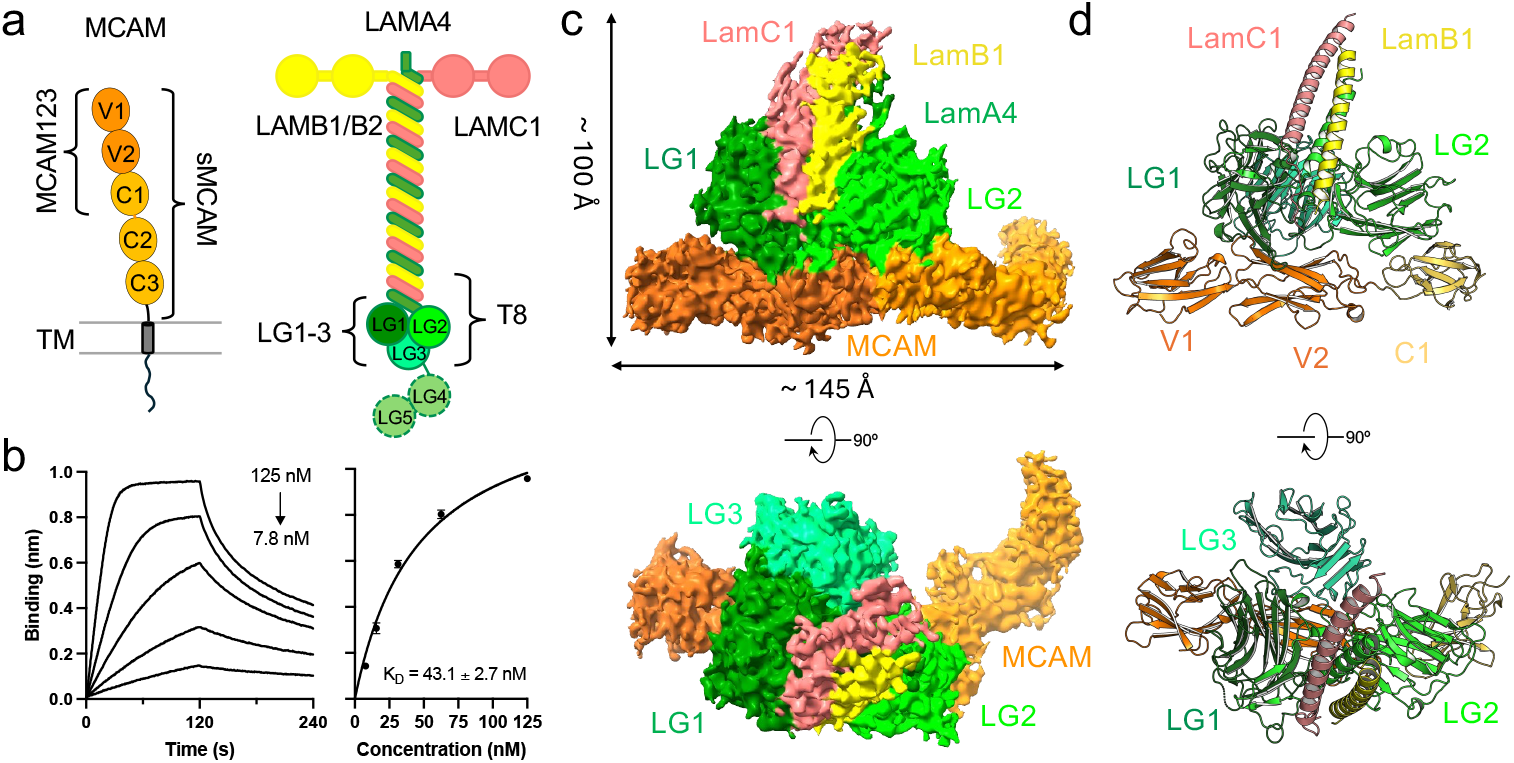
MCAM interacts with T8-LM411. **a**, Architecture scheme of MCAM and LamA4. The expression constructs used in this study are labeled by the side of curly brackets. **b**, Direct binding of T8-LM411to sMCAM detected by biolayer interferometry (BLI). The sensors were loaded with sMCAM. The binding affinity represents mean ± SEM from three independent experiment. **c**, Surface view of the Cryo-EM reconstruction of sMCAM-T8-LM411 complex (4.13 Å). **d**, Ribbon diagram of an atomic model of sMCAM-T8-LM411 complex. MCAM C2-C3 domains are not modeled as the resolution is over 4.4 Å (Supplementary Fig. 2).

Biolayer interferometry (BLI) revealed that sMCAM binds T8-LM411 (LamA4: N757–E1405; laminin β1: D1714–L1786; laminin γ1: D1528–P1609) with an equilibrium dissociation constant (KD) of 43 nM (**Fig. 1b**), representing an approximately 40-fold stronger affinity compared to its interaction with LamA4 LG1-3 (residues Q833–E1405) (**Supplementary Fig. 1b**). This is consistent with prior observations that the laminin β and γ chains promote a more compact assembly by stabilizing the packing of LG domains[22], which may enhance MCAM binding affinity in solution.

For structural characterization, either T8-LM411 or LamA4 LG1-3 was mixed with sMCAM at a 1:1 molar ratio and subjected to cryo-EM (**Supplementary Fig. 2 and 3**). Image processing using cryoSPARC[23, 24] yielded a reconstruction at 4.13 Å resolution for sMCAM-T8-LM411 complex (**Fig. 1c and d, Supplementary Fig. 2, Supplementary Table 1**). Although the resolution is reduced due to flexibility in the laminin coiled-coil and distal sMCAM domains, the core involving MCAM and LamA4 interactions reaches a resolution better than 3.6 Å (**Supplementary Fig. 2f**). We also determined the cryo-EM structure of sMCAM in complex with LamA4 LG1-3 at 3.22 Å resolution (**Fig. 2a and b, Supplementary Fig. 3, Supplementary Table 1**). Both structures show the same overall interaction. The complex adopts a compact architecture in which LamA4 LG1-3 forms a cloverleaf-like structure reminiscent of integrin-binding laminin fragments[22], while MCAM wraps along its surface. Two distinct interaction sites are observed. Site 1 is formed between the first globular domain of LamA4 (LG1) and the second Ig domain of MCAM (V2). Here, a series of main-chain interactions is made between MCAM (residues Q191–S198) and the V2 domain and LamA4 (residues T955–I961). Site 2 is formed between the second globular domain of LamA4 (LG2) and the third Ig domain of MCAM (C1). This interaction is dominated by hydrophobic contacts between MCAM’s L323 and LamA4 residues I1142 and I1150 (**Fig. 2b, Supplementary Fig. 4a and b**).

**Figure 2.**
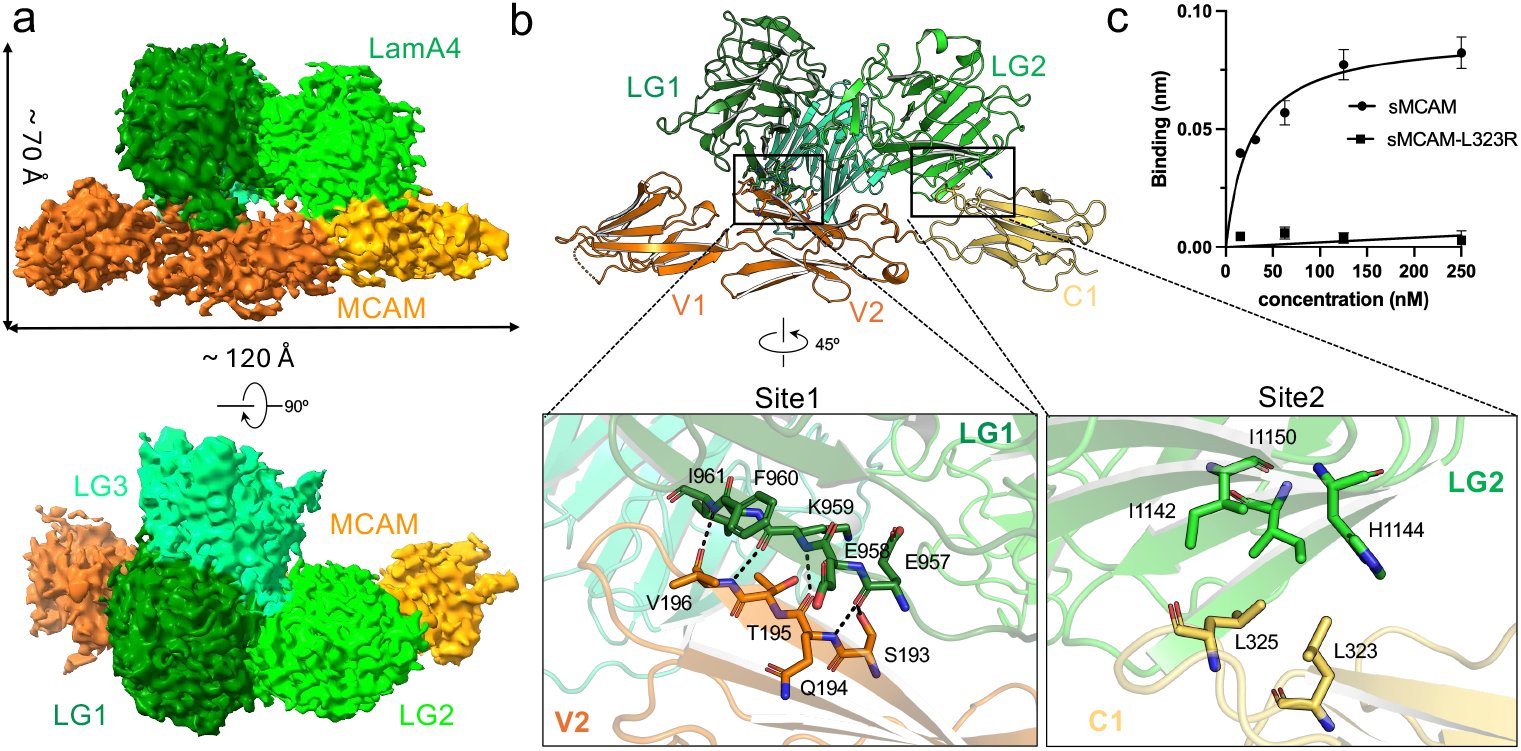
MCAM engages with LamA4 LG1-3 at two distinct sites. **a**, Surface view of the Cryo-EM reconstruction of the complex of sMCAM with isolated LamA4 LG1-3 domains (3.22Å). **b**, Enlarged detailed view of site 1 interactions between MCAM V2 domain and LamA4 LG1 domain, and site 2 interactions between MCAM C1 domain and LamA4 LG2 domain. **c**, Binding of MCAM L323R mutant to T8-LM411 measured by BLI. The sensors were loaded with T8-LM411. The binding affinity represents mean ±SEM from three independent experiments.

Mutational analysis supports the structural observations. Substitution of MCAM L323 with arginine abolishes binding to LamA4, while analogous mutations in LamA4 (I1142R or I1150R) weaken the binding affinities by about 5.9 fold for the I1142R mutant, and 2.7 fold for the I1150R mutant (**Fig. 2b, Supplementary Fig. 4c**). These results establish a bipartite binding mode in which hydrophobic interactions at site 2 are critical for complex stability.

Our structure indicates that MCAM engages exclusively with the laminin α4 chain. To investigate whether MCAM binds LM411 and LM421 differently, we measured the binding of T8-LM421 (laminin β2: A1727–Q1798) to MCAM on BLI. The binding curves show a similar affinity (KD = 41.5 nM) to that observed for T8-LM411 (**Fig. 3a)**. We further determined a cryo-EM structure of MCAM in complex with T8-LM421 at 3.76 Å resolution (**Fig. 3b and c, Supplementary Fig. 5)**. Similar to MCAM-T8-LM411, the laminin coiled-coil and distal MCAM domains of MCAM-T8-LM421 are flexible, but the core involving MCAM V2-C1 and LamA4 LG1-2 interactions is stable, with a resolution better than 3.3 Å (**Supplementary Fig. 5e**). Comparison of MCAM-LM411 and MCAM-LM421 complexes shows nearly identical binding interfaces, demonstrating that MCAM specifically recognizes the LG1 and LG2 domains of LamA4 and does not discriminate between the b1 and b2-containing isoforms LM411 and LM421.

**Figure 3.**
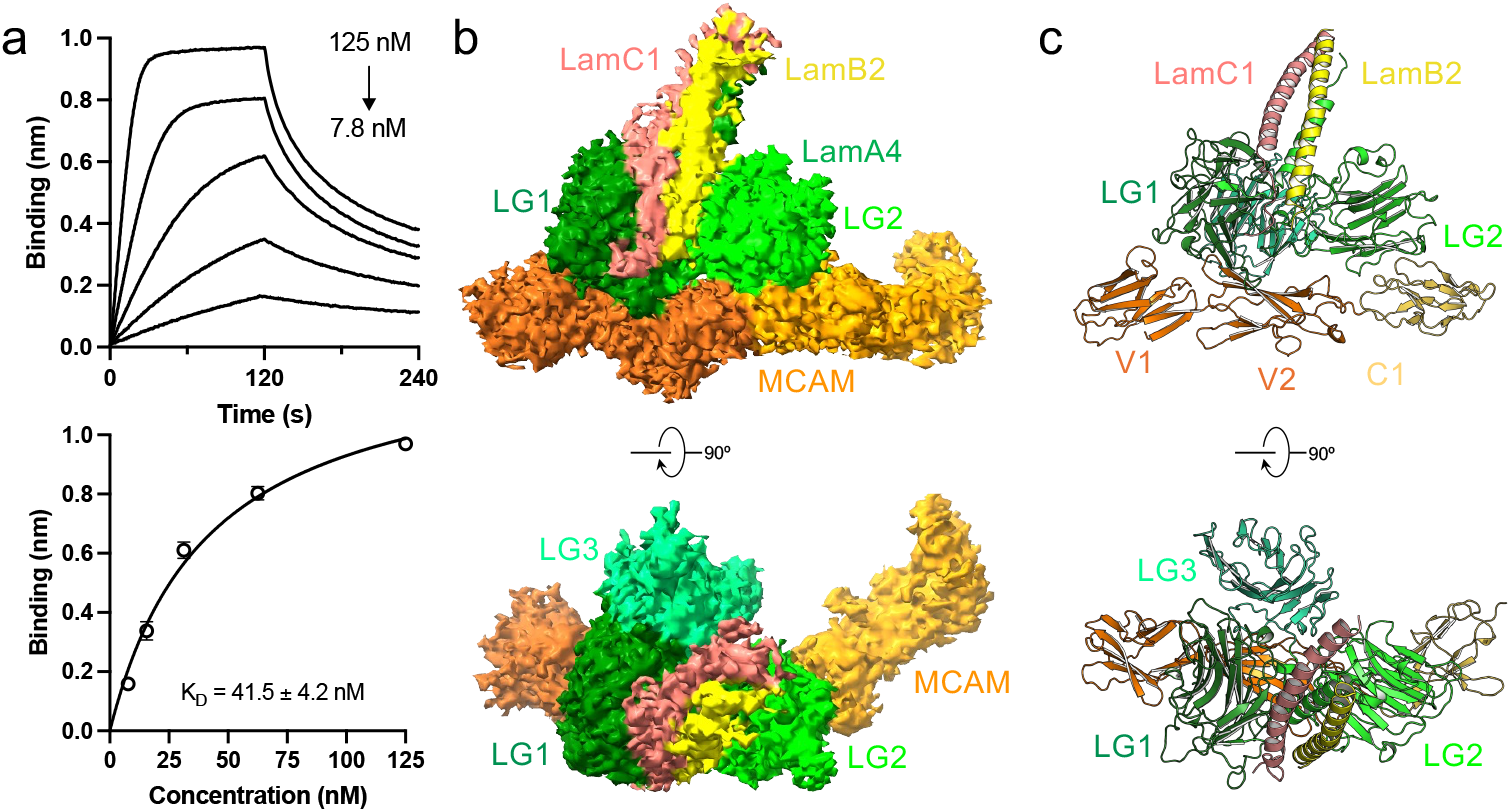
MCAM interacts with T8-LM421. **a**, Direct binding of T8-LM421 to sMCAM detected by BLI. The sensors were loaded with sMCAM. The binding affinity represents mean ±SEM from three independent experiments. **b**, Surface view of the Cryo-EM reconstruction of sMCAM-T8-LM421 complex (3.76Å). **c**, Ribbon diagram of an atomic model of sMCAM-T8-LM421 complex.

### MCAM and Integrin α6β1 share overlapping binding sites on Laminin α4

A prior structure of Integrin α6β1 bound to Laminin α5 (LamA5) revealed how integrins engage the laminin α-chain globular domains. However, there remains no experimental structure showing how Integrin α6β1 binds specifically to LamA4. Therefore, to make a direct comparison to our structure, we used structural modeling with AlphaFold 3[25] to predict the integrin α6β1 – T8-LM411 complex. The predicted complex is of high confidence (**Supplementary Fig.6a, ipTM = 0.79**), and uses the same interface as the experimental and highly similar LamA5 complex[10]. The α6β1-LamA4 interaction is mediated by residues that coincide with the MCAM-binding interface. Specifically, the LG1 β-sheet region (E957–K963; site 1) and hydrophobic residues in LG2 (including I1142 and I1150; site 2) are implicated in both MCAM and integrin binding (**Fig. 4a and b, Supplementary Fig. 6a**). Isoform-specific differences further support this model. Integrin α6×1β1 and α7×1β1 lack key elements required for site 1 interaction, suggesting partial overlap, whereas integrin α7×2β1 shows limited compatibility with the LamA4 interface, consistent with prior reports of weak or absent binding[9, 26, 27] (**Supplementary Fig. 6b-d**).

**Figure 4.**
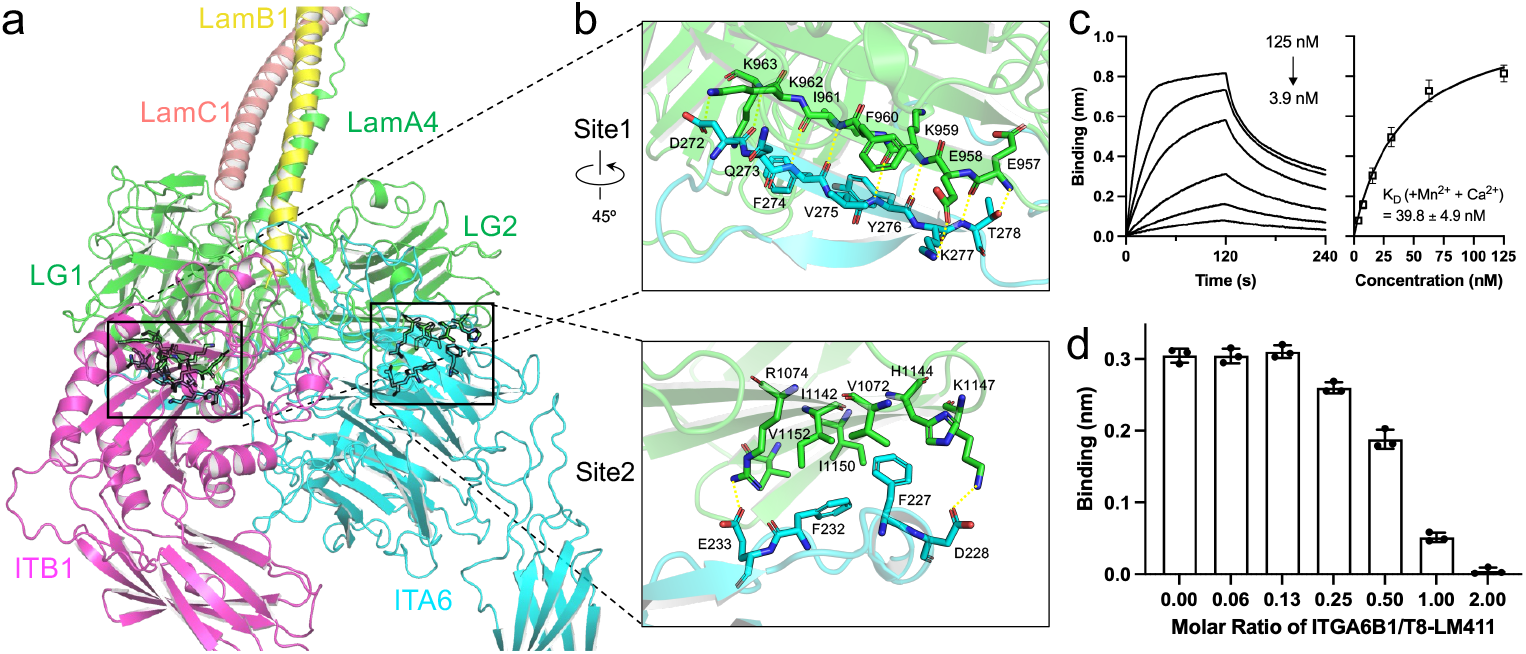
MCAM and Integrin α6β1 share overlapping binding sites on Laminin α4. **a**, AlphaFold 3 predicted complex model of T8-LM411 and integrin α6β1. The pLDDT and ipTM of the prediction is shown in Supplementary Fig. 5. **b**, Enlarged detailed view of site 1 interactions between integrin α6 and LamA4 LG1 domain, and site 2 interactions between integrin α6 and LamA4 LG2 domain. **c**, Binding of T8-LM411 to sMCAM in the buffer containing divalent metal cations detected by BLI. The sensors were loaded with sMCAM. The binding affinity represents mean ±SEM from three independent experiments. **d**, Competition between integrin α6β1 and sMCAM for LM411 binding measured by BLI. The sensors were loaded with sMCAM. 20 nM of T8-LM411 was mixed with increased concentrations of integrin α6β1. The responses at the plateau were plotted against the molar ratio of integrin α6β1 over T8-LM411. Bars represent mean ±SD from three independent experiments.

We experimentally tested this competition using BLI. Since integrin-laminin interactions are divalent metal cation-dependent[9, 10], we first confirmed that the presence of Mn2+ and Ca2+ does not affect MCAM–LamA4 binding (**Fig. 4c**). We then assessed whether integrin α6β1 could compete with MCAM for LM411 binding. Increasing concentrations of integrin α6β1 (ITGA6B1) progressively reduced T8-LM411’s ability to associate with sensor bound MCAM, demonstrating direct competition for overlapping binding sites (**Fig. 4d**). At a 2:1 ratio of ITGA6B1 to T8-LM411, no MCAM binding was detected, indicating that all MCAM binding sites were prebound with integrin.

### MCAM inhibits Laminin α4-mediated melanoma cell migration

To evaluate the functional consequences of the MCAM-LamA4 interaction, we performed migration assays using MeWo melanoma cells. These cells exhibit robust migration towards both LM411 and LM421, with LM421 showing greater potency at comparable or lower concentrations (**Fig. 5a, b**), consistent with previous reports[19]. In contrast, isolated LamA4 LG domains failed to induce migration, indicating that intact laminin heterotrimers are required for this process (**Fig. 5a**). Given that MeWo cells express low levels of MCAM (**Supplementary Fig. 7a**), the observed migration is likely mediated by integrins rather than by MCAM. To exclude a major contribution from MCAM expressed on the cell surface, we performed the same assay using MCF7 breast cancer cells, which express low levels of both ITGA6 and ITGB1 compared to MeWo cells (https://www.proteinatlas.org/). MCF7 cells show little to no migration toward LM411 or LM421. Ectopic expression of MCAM in MCF7 cells does not enhance migration (**Supplementary Fig. 7a and b**), further demonstrating that laminin-dependent migration is driven by integrins and not MCAM.

**Figure 5.**
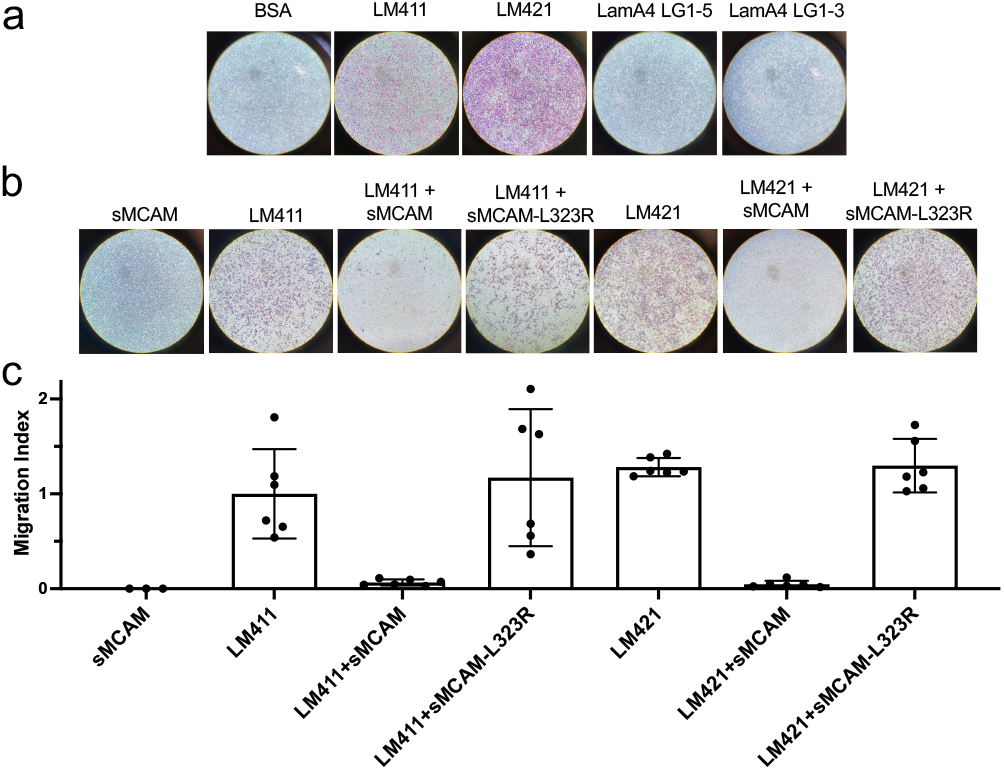
MCAM inhibits Laminin α4-mediated melanoma cell migration. **a**, LM411 and LM421 drive MeWo cell migration but not LamA4 LG domains. The concentration of the laminin proteins is 250 nM. **b**, sMCAM inhibits MeWo cell migration towards LM411 and LM421. The binding-deficient sMCAM-L323R mutant failed to inhibit the migration. The concentrations of LM411 and LM421 are 100 nM and 25 nM, respectively. The concentration of sMCAM and sMCAM-L323R is 1 μM. **c**, Quantification of the migration assays shown in **b**. Two transwell inserts were analyzed per condition, and each experiment was repeated three times for statistical analysis.

Based on the shared binding interface, we hypothesized that MCAM could competitively inhibit integrin-dependent migration. Indeed, addition of sMCAM with LM411 or LM421 effectively blocked MeWo migration toward LM411 and LM421 (**Fig. 5b and c**). The inhibitory effect was recapitulated by the V1-C1 fragment of MCAM (MCAM123), indicating that the N-terminal domains are sufficient for competition (**Supplementary Fig. 7d and e**). In contrast, an in-house-generated Fab, which recognizes the LamA4 LG2 domain at an epitope distinct from the MCAM/integrin binding site (**Supplementary Fig. 7c and 8**), did not block MeWo migration (**Supplementary Fig. 7d and e**). Moreover, the binding-deficient MCAM L323R mutant failed to inhibit migration, confirming that blockade depends on direct interaction with LamA4 (**Fig. 5b and c**). These results demonstrate that MCAM functions as a competitive inhibitor of integrin-mediated migration by occupying shared binding sites on laminin α4.

## Discussion

Here we show the structural basis for MCAM recognition of laminin α4 and demonstrate that MCAM directly competes with integrin α6β1 for a shared laminin interface. Previous work established that α4-containing laminins, particularly laminin-411 and laminin-421, support cell migration through α6β1 integrin and MCAM/CD146-dependent mechanisms[15, 19]. Antibody-mapping experiments further suggested that α6β1 integrin and MCAM bind at closely apposed sites within the laminin α4 globular domain[20]. However, without a high-resolution structure of the MCAM-laminin complex, it remained unclear whether these receptors could bind laminin cooperatively or instead compete for the same physical surface. Our cryo-EM structure resolves this question by showing that MCAM engages a surface centered on the laminin α4 LG1-LG2 domains and occupies the region used by laminin-binding integrins.

Laminin-binding integrins recognize a composite T8 surface whose activity depends on the α-chain LG domains together with β- and γ-chain contributions, including the γ-chain C-terminal tail. This architecture allows integrins to interpret the broader laminin heterotrimer code, with different integrin heterodimers displaying overlapping but distinct preferences across laminin isoforms. By contrast, MCAM specifically targets the laminin α4 globular region with relative independence from β/γ-chain identity. This mode of recognition is well suited to act specifically at α4-laminin-rich environments, including vascular and perivascular basement membranes, post-capillary venules, lymphatic and tumor-associated vessels, and inflammatory barrier-crossing sites such as the neurovascular unit. In these settings, MCAM may function as a directed α4-laminin reader. Thus, MCAM may provide a mechanism for selectively engaging α4-rich vascular or stromal microenvironments while bypassing some of the β/γ-chain constraints that govern integrin binding. At the same time, because MCAM occupies a laminin α4 surface shared by integrins, MCAM binding could either promote localization to α4-laminin-rich niches or, when present in soluble or matrix-bound form, mask integrin-accessible sites and restrain integrin-driven migration.

These findings have therapeutic implications. Integrins are attractive but challenging drug targets because they are widely expressed, functionally pleiotropic and involved in normal adhesion, immunity, hemostasis and tissue homeostasis[28, 29]. Direct systemic integrin blockade can therefore produce broad and unwanted effects. The MCAM-laminin α4 structure identifies an alternative strategy: targeting the laminin α4 receptor-binding surface itself. Biologics that mimic the MCAM footprint, stabilize an MCAM-like blocked state, or use engineered MCAM-derived scaffolds could selectively prevent α6β1 integrin engagement on α4-containing laminins without globally inhibiting integrin function. Because laminin α4-containing vascular basement membranes regulate leukocyte trafficking and tumor-cell migration[6, 7, 15, 19, 20], such approaches may be relevant to metastatic dissemination, neuroinflammation and inflammatory cell trafficking.

Several limitations should be considered. The current work defines a competitive mechanism using purified proteins, structural analysis and in vitro migration assays. It does not directly quantify receptor-density thresholds, cell-surface occupancy, soluble versus membrane-tethered MCAM balance, or in vivo basement membrane occupancy. It also does not exclude additional MCAM functions mediated by the cytoplasmic tail, lateral membrane organization, endothelial interactions or soluble ectodomain signaling[16-18, 30]. Rather, the study defines a precise structural mechanism by which MCAM can inhibit laminin-dependent migration: direct competition with α6β1 integrin for laminin α4.

In summary, we show that MCAM and integrin α6β1 compete for a shared laminin α4 interface. This high-resolution structure explains how MCAM occupancy can suppress integrin-dependent migration, resolves a long-standing ambiguity in MCAM-laminin biology, and establishes a structural foundation for therapeutic strategies aimed at selectively blocking pathological cell migration.

## Acknowledgements

We thank the Yale Cancer Biology Institute and its laboratories for valuable discussions. We thank Jianfeng Lin at the Yale Cryo-EM Resource Center for assistance in Cryo-EM sample screening and data collection. We thank Yuhong Zuo and Long Han for their expertise in guiding the CryoEM processing. Yale Cryo-EM Resource is funded in part by the NIH grant S10OD023603. This work was supported by the NIH grants RM1GM149406 (D.E.K.).

## Author contributions

T.L. and D.E.K. wrote the manuscript with input from C.R.A.. T.L. generated all materials, performed and analyzed all the experiments assisted by S.A. and H.L.

## Materials and Methods

### Recombinant Protein Expression and Purification

Codon-optimized cDNAs encoding human MCAM (V24 – H646), laminin α4 (LamA4, N757 – E1405), laminin β1 (LamB1, D1714 – L1786), laminin β2 (LamB2, A1727 – Q1798), and laminin γ1 (LamC1, D1528 – P1609) were synthesized by IDT. The MCAM and LamA4 constructs were cloned into mammalian expression vectors containing a Wnt3 signal peptide (MEPHLLGLLLGLLLGGTRVLAG) and an N-terminal octa-histidine tag followed by a Factor Xa cleavage site (GHHHHHHHHGSSTSNGTIEGRS). The LamB1 and LamB2 constructs were cloned into a mammalian expression vector containing an IgG κ-chain signal peptide (MEFQTQVLMSLLLCMSGAAA). The LamC1 construct was cloned into mammalian expression vector containing an IgG γ-chain signal peptide and an N-terminal twin-strep tag (MKLPVLLVVLLLFTSPASSSSAWSHPQFEKGGGSGGGSGGSAW-SHPQFEK). MCAM and LamA4 LG truncations and mutations were conducted by In-Fusion Snap Assembly (Takara). Full-length LM411 and LM421 were purchased from Biolamina (cat# LN411-02, LN421-02) or Acro Biosystems (cat# LAT-H5263-100ug). Integrin α6β1 was purchased from Acro Biosystems (cat# IT1-H52W7-100ug). LamA4 LG1-5 was purchased from R&D systems (cat# 7340-A4).

Recombinant proteins were expressed by transient transfection of Expi293 cells (Thermo Fisher Scientific). Conditioned medium was collected 3-4 days post-transfection and applied directly to Ni-Penta™ Agarose-Base Resins (Marvelgent Biosciences) by gravity flow. The resin was washed with buffer containing 15 mM imidazole, 100 mM NaCl, pH 8.0, and bound proteins were eluted with buffer containing 300 mM imidazole, 100 mM NaCl, pH 8.0. For further purification, eluates were subjected to size-exclusion chromatography on a HiLoad 26/600 Superdex 200 pg column or a Superdex 200 Increase 10/300 GL size exclusion column (Cytiva Life Sciences), equilibrated in 20 mM HEPES, 100 mM NaCl, pH 7.5. Peak fractions were analyzed by SDS-PAGE, concentrated using 10 kDa or 30 kDa Centricon spin concentrator (Millipore), and stored at 4 ^°^C.

### Cryo-EM structure determination

The MCAM-LamA4 complex was prepared by mixing purified MCAMecd and LamA4 LG1-3 at a 1:1 molar ratio and diluting the mixture to approximately 0.3 mg/ml. Three microliters of sample were applied to glow-discharged Quantifoil holey carbon grids (R1.2/1.3 300 mesh). Grids were blotted for 6 s at 16 ^°^C with 100 % humidity before plunge-freezing into liquid ethane using a Vitrobot Mark IV (FEI). Cryo-EM data were collected on a Glacios microscope equipped with a K3 summit direct electron detector (Gatan) using SerialEM software in super-resolution mode at a nominal magnification of 45,000×, corresponding to a calibrated super-resolution pixel size of 0.434 Å. Micrographs were acquired over a defocus range of -0.8 to -2.0 µm. The dose rate was set to 6.3 electrons per physical pixel per second. The exposure time of each movie was about 1.2 s, leading to a total dose of approximately 40 electrons per Å^2^, fractionated into 40 frames. A total of 7,387 images were collected for MCAM-LamA4 complex.

Image processing was performed in cryoSPARC[24] v4.3.018 following the standard workflows. During curation, micrographs with CTF-estimated resolution worse than 6 Å, average intensity greater than 50, or relative ice thickness greater than 1.2 Å were excluded. Initial particle picking was performed using blob picking, followed by several rounds of 2D classification. Particles belonging to good 2D classes were used for ab initio reconstruction to generate initial 3D volumes. Particles corresponding to the complex were then selected for Topaz[31]-based particle picking. The resulting particle set was subjected to iterative 2D classification, ab initio reconstruction, and heterogeneous refinement to obtain a high-quality particle class for final map generation. The best map was refined by non-uniform refinement[23], and mask corresponding to the compact MCAM and LamA4 LG1-3 regions were generated by volume segmentation in ChimeraX v1.11.1 followed by local refinement. Overall resolution was estimated using the gold-standard Fourier shell correlation (FSC) =0.143 criteria and local resolution was calculated in cryoSPARC from the two half-maps.

### Model building and refinement

An initial model was built into the cryo-EM density using the Phenix Predict and Build: Cryo-EM module (version 1.21rc1-5127)[32]. The model was refined iteratively using Phenix Real-space Refinement module[33] and Coot[34]. Detailed refinement statistics and model-validation metrics are provided in **Supplementary Data Table 1**.

### AlphaFold models

The AlphaFold 3 Server was used to generate a model of the truncated E8-LM411-integrin α6β1 complex using residues N757 – E1405 of LAMA4 (Uniprot ID: Q16363), D1714 – L1786 of LAMB1 (Uniprot ID: P07942), D1528 – P1609 of LAMC1 (Uniprot ID: P11047), F24 – K680 of ITGA6 (Uniprot ID: P23229), and Q21 – E465 of IGB1 (Uniprot ID: P055556).

### Bio-layer Interferometry

Bio-layer Interferometry (BLI) was performed using an Octet RED96e instrument (Sartorius). ProteinA biosensors (Sartorius) were loaded with Fc fused MCAMecd to a response level of approximately 1 nm, equilibrated in working buffer (20 mM HEPES, 100 mM NaCl, pH 7.5) for 30 s, and then immersed in wells containing the indicated concentrations of T8-LM411, T8-LM421, or mutant proteins for 120 s, followed by dissociation in working buffer for 120 s. Equilibrium (plateau) responses were plotted against concentration using GraphPad Prism 10.3.1. For competition assay, 20 nM T8-LM411 was mixed with increasing concentrations of integrin α6β1 in working buffer supplemented with 1 mM MnCl_2_, 0.1 mM CaCl_2_.

### Transwell Migration Assay

Transwell migration assays were performed essentially as described previously[19]. Briefly, the underside of transwell inserts (Corning cat # 3422) was coated with LM411 (Biolamina or Acro Biosystems) or LM421 (Biolamina or Acro Biosystems) or with laminin pre-complexed with 1 μM MCAM at 37 ^°^C for 2 h, then blocked with DMEM containing 0.1% BSA for 30 min. Cells were detached using Cellstripper (Corning cat# 25-056-CI) and resuspended in DMEM supplemented with 0.1% BSA medium. A total of 1 × 10^5^ cells were added to the upper chamber, and DMEM with 0.1% BSA was added to the lower chamber. After 18 hours at 37 °C, transwell inserts were washed six times with sterile PBS, fixed with 4% paraformaldehyde (Santa Cruz cat# sc-281692) for 15 min, and stained using a hematoxylin and eosin staining kit (Abcam cat# ab245880) according to the manufacturer’s instructions. Two transwell inserts were analyzed per condition, and each experiment was repeated at least three times for statistical analysis.

### Western Blotting

Cells were washed with PBS and lysed in buffer containing 20mM Tris-HCl pH 8.0, 150mM NaCl, and 0.5% NP-40, supplemented with protease inhibitor cocktail (Cell Signaling Technology). Lysates were incubated on ice for 30 min and clarified by centrifugation at 15,000 rpm for 15 min. Protein concentrations were determined using a Bradford assay (Thermo Fisher Scientific). For detection of MCAM, integrin α6, and integrin β1, 50 μg total protein per sample was separated on 4-20% gradient gel for SDS-PAGE gels using BLUEstain 2 protein ladder (GoldBio) as a molecular-weight ladder. Proteins were transferred to PVDF membrane (EMD millipore) in transfer buffer containing 25 mM Tris, 192 mM glycine, 20% methonal (v/v) using a constant voltage of 90 V for 45 min. Membranes were blocked with 5% non-fat milk (Americo) in TBST buffer (20 mM Tris-HCl pH8.0, 150 mM NaCl, 0.1% Tween 20) for 1 h at room temperature followed by incubating with the primary antibodies for 1h at room temperature. Primary antibodies were used at the following dilutions: MCAM (Cell Signaling Technology cat# 81701), 1:1000; integrin α6 (Cell Signaling Technology cat# 3750), 1:1000; integrin β1 (Cell Signaling Technology cat# 9699T, 1:1000). Membranes were then incubated with horseradish peroxidase (HRP)-conjugated goat anti-rabbit IgG secondary antibody (Cell Signaling Technology) at 1:5000 dilution for 1 h at room temperature. The membranes were developed with Amersham ECL prime western blotting detection reagent (Cytiva Life Sciences) and chemiluminescence scanned using ChemiDoc (Bio-Rad).

**Supplementary Figure S1.**
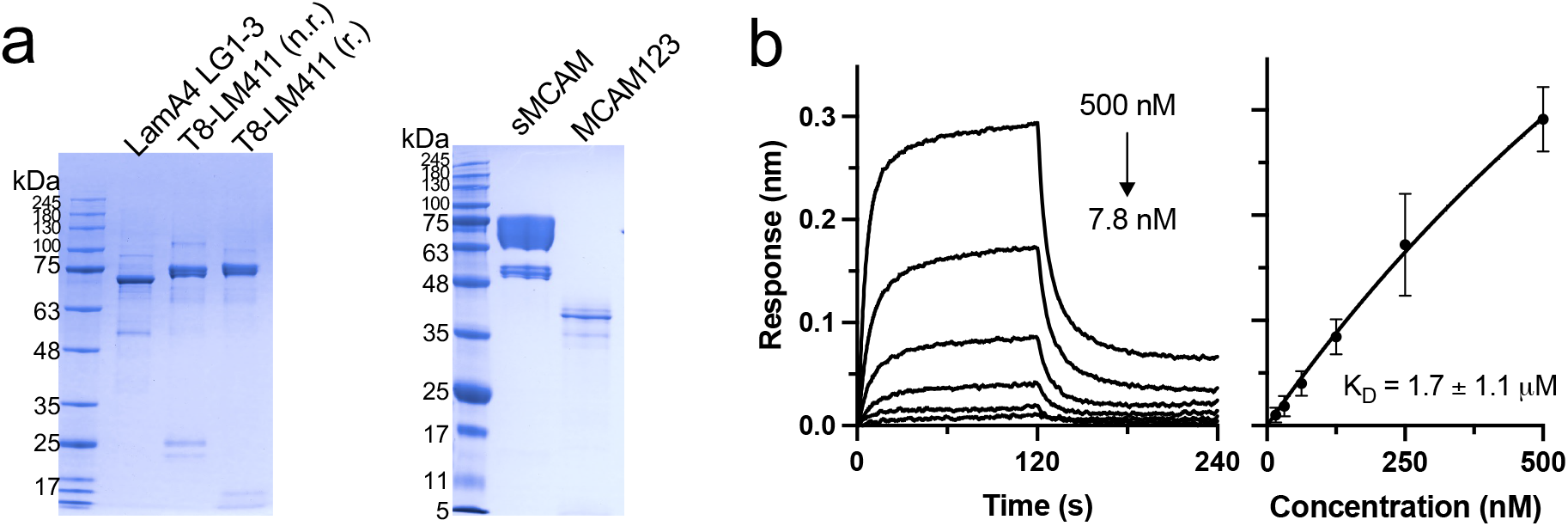
Recombinant LamA4 and MCAM proteins. **a**, SDS-PAGE gel stained with Coomassie blue shows the sizes of different constructs of LamA4 and MCAM expressed. **b**, Direct binding of LamA4 LG1-3 to sMCAM detected by biolayer interferometry (BLI). The sensors were loaded with sMCAM. The binding affinity represents mean ± SEM from three independent experiments.

**Supplementary Fig 2.**
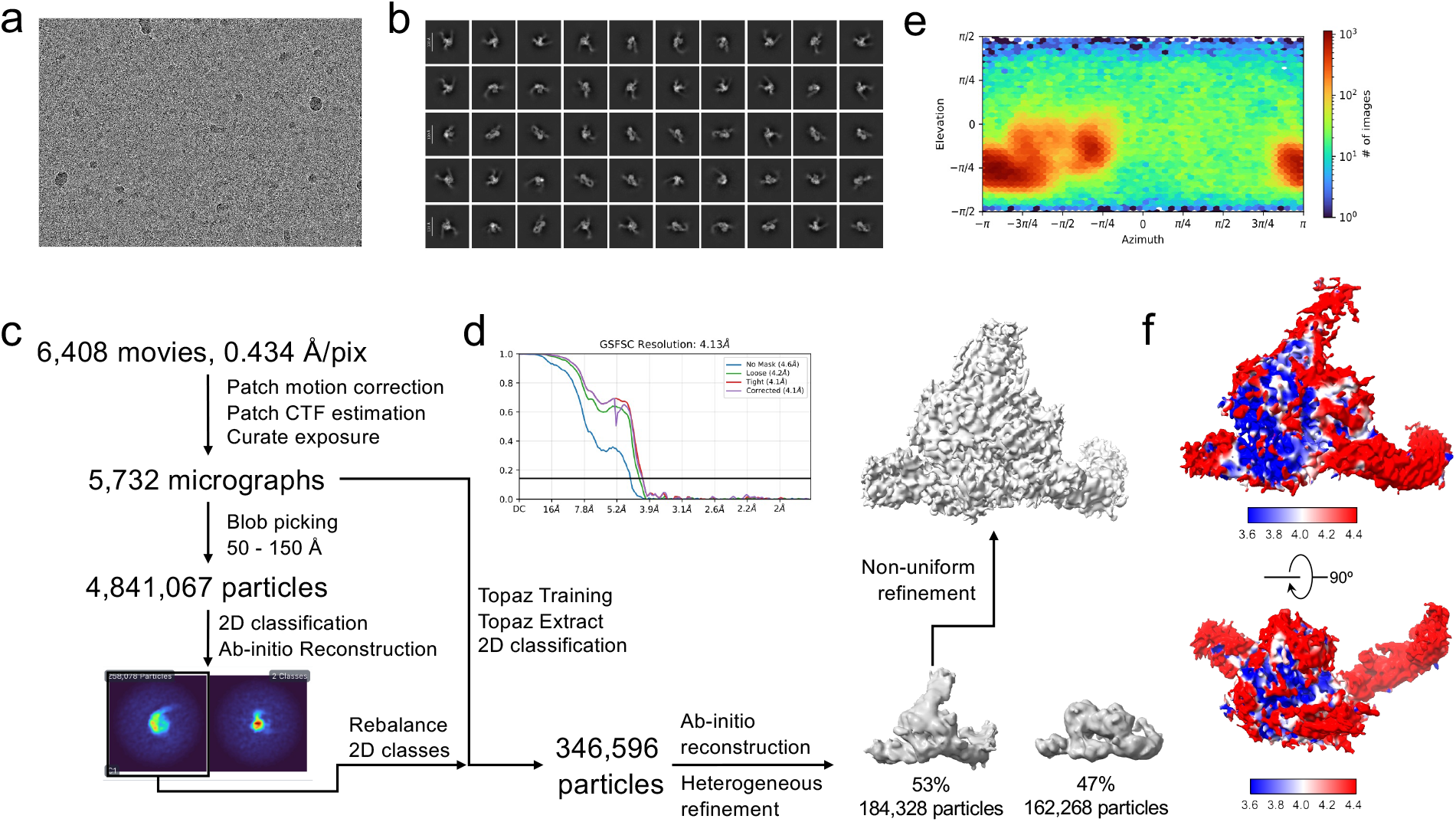
Cryo-EM data processing for the MCAM-T8-LM411 complex structure. **a**, Representative raw micrograph of sMCAM-T8-LM411 complex. **b**, Representative 2D class averages of sMCAM-T8-LM411 complex. **c**, Images of the processing workflow for structure determination of sMCAM-T8-LM411 complex. **d**, Gold-standard Fourier Shell Correlation (FSC) of the local-refined map. **e**, Angular distribution of particles in the final map obtained by non-uniform refinement. **f**, Local resolution estimation of the map.

**Supplementary Fig 3.**
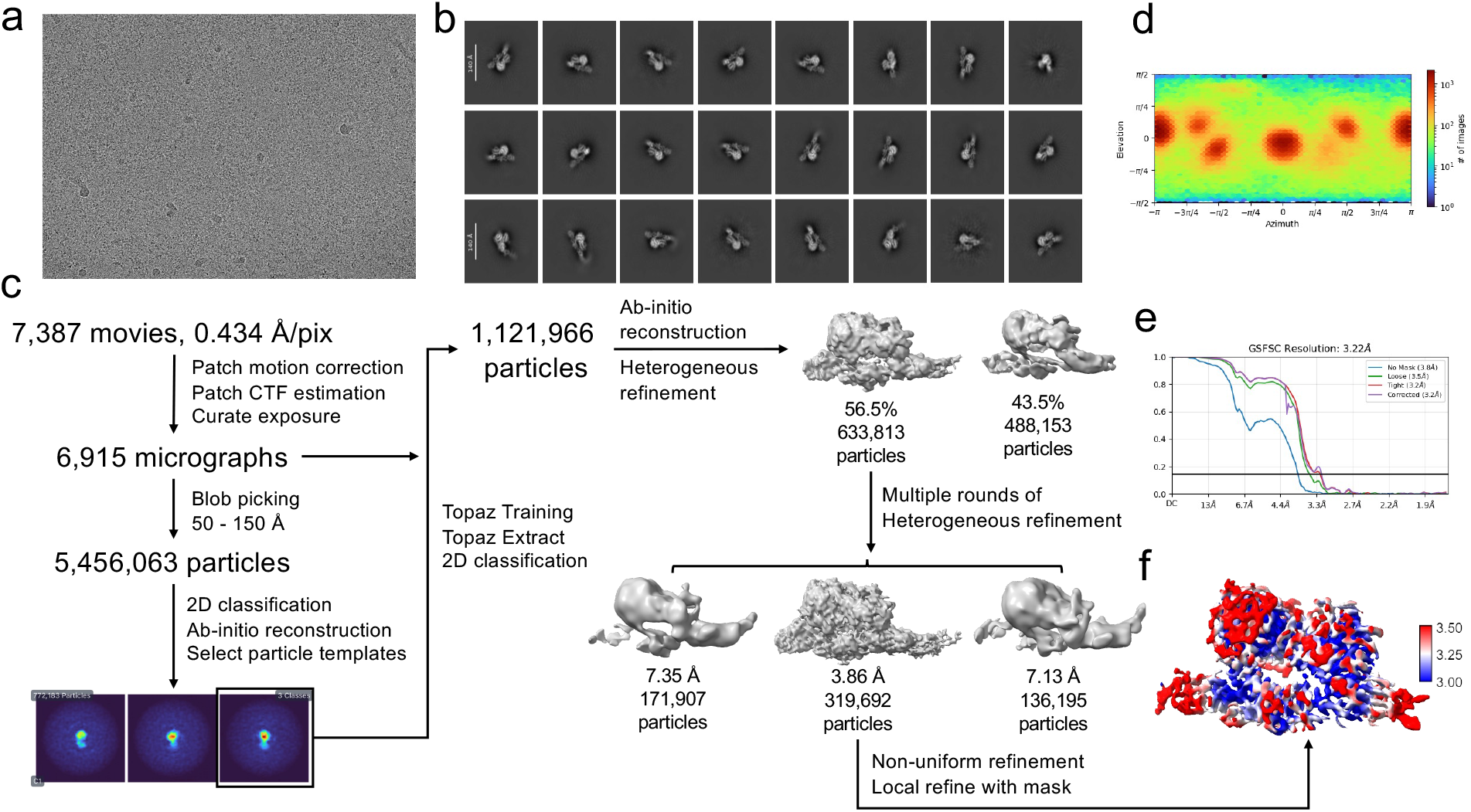
Cryo-EM data processing for the MCAM-LamA4 LG1-3 complex structure. **a**, Representative raw micrograph of sMCAM-LamA4 LG1-3 complex. **b**, Representative 2D class averages of sMCAM-LamA4 LG1-3 complex. **c**, Images of the processing workflow for structure determination of sMCAM-LamA4 LG1-3 complex. **d**, Angular distribution of particles in the final map obtained by non-uniform refinement. **e**, Gold-standard Fourier Shell Correlation (FSC) of the local-refined map. **f**, Local resolution estimation of the map.

**Supplementary Fig 4.**
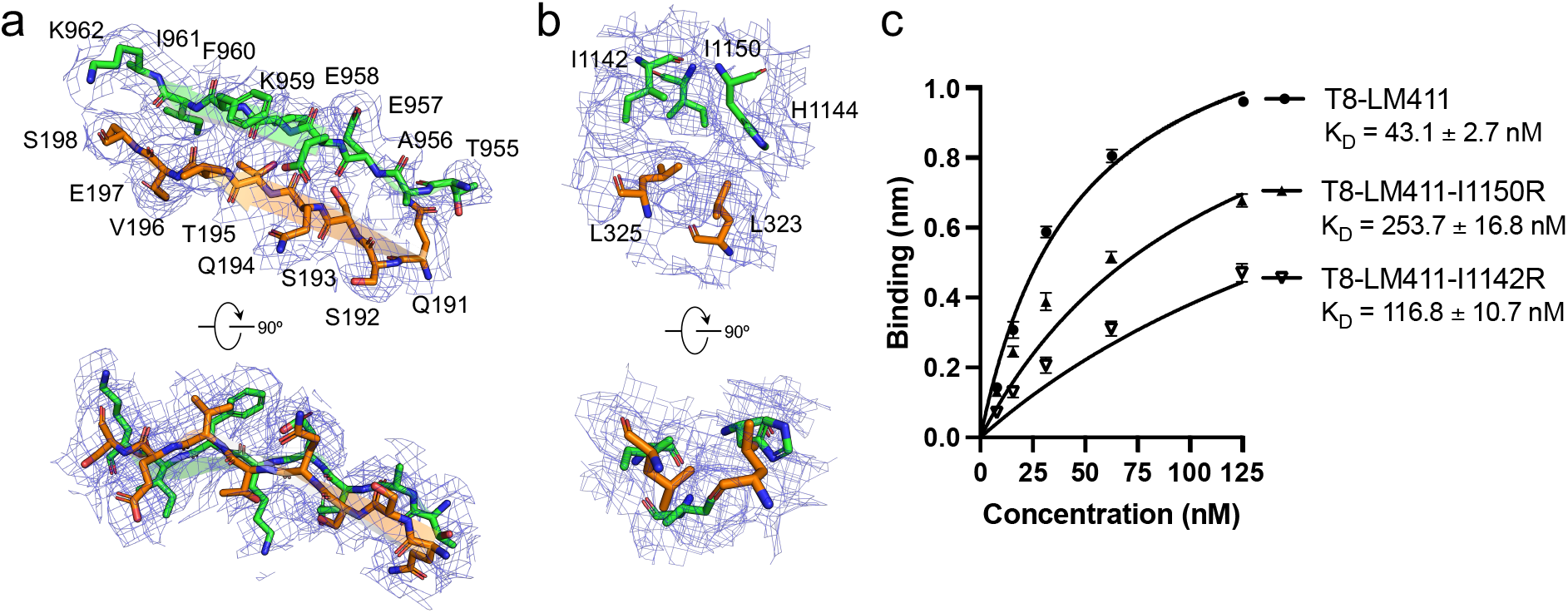
Regional density map of MCAM-LamA4 interaction sites. **a**, Density map of MCAM-LamA4 interaction site1. **b**, Density map of MCAM-LamA4 interaction site2. **c**, Binding of T8-LM411 mutants to MCAM detected by biolayer interferometry (BLI). The sensors were loaded with sMCAM. The binding affinity represents mean ± SEM from three independent experiments.

**Supplementary Fig 5.**
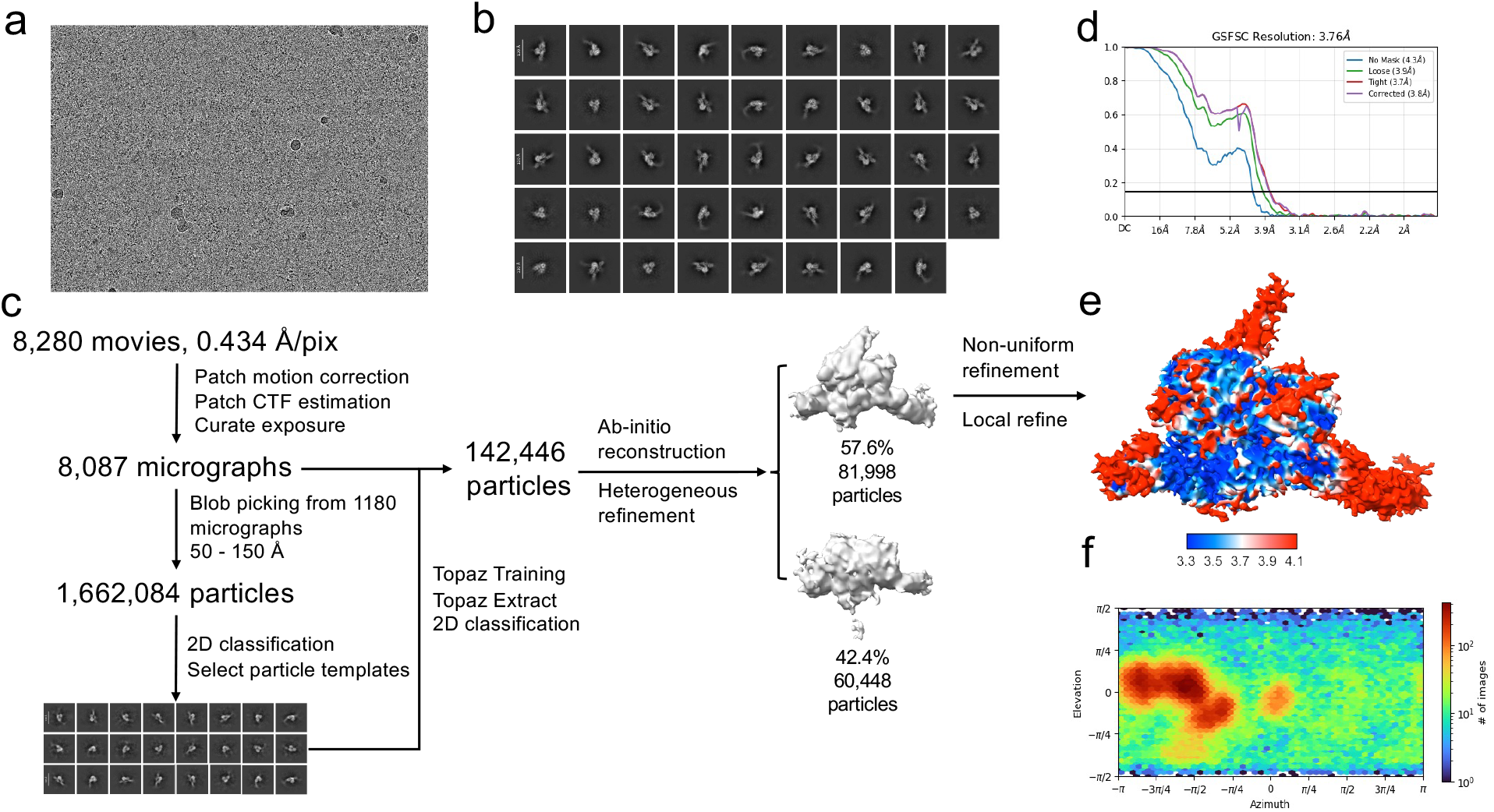
Cryo-EM data processing for the MCAM-T8-LM421 complex structure. **a**, Representative raw micrograph of sMCAM-T8-LM421 complex. **b**, Representative 2D class averages of sMCAM-T8-LM421 complex. **c**, Images of the processing workflow for structure determination of MCAM-T8-LM421 complex. **d**, Gold-standard Fourier Shell Correlation (FSC) of the local-refined map. **e**, Local resolution estimation of the map. f, Angular distribution of particles in the final map obtained by non-uniform refinement.

**Supplementary Fig 6.**
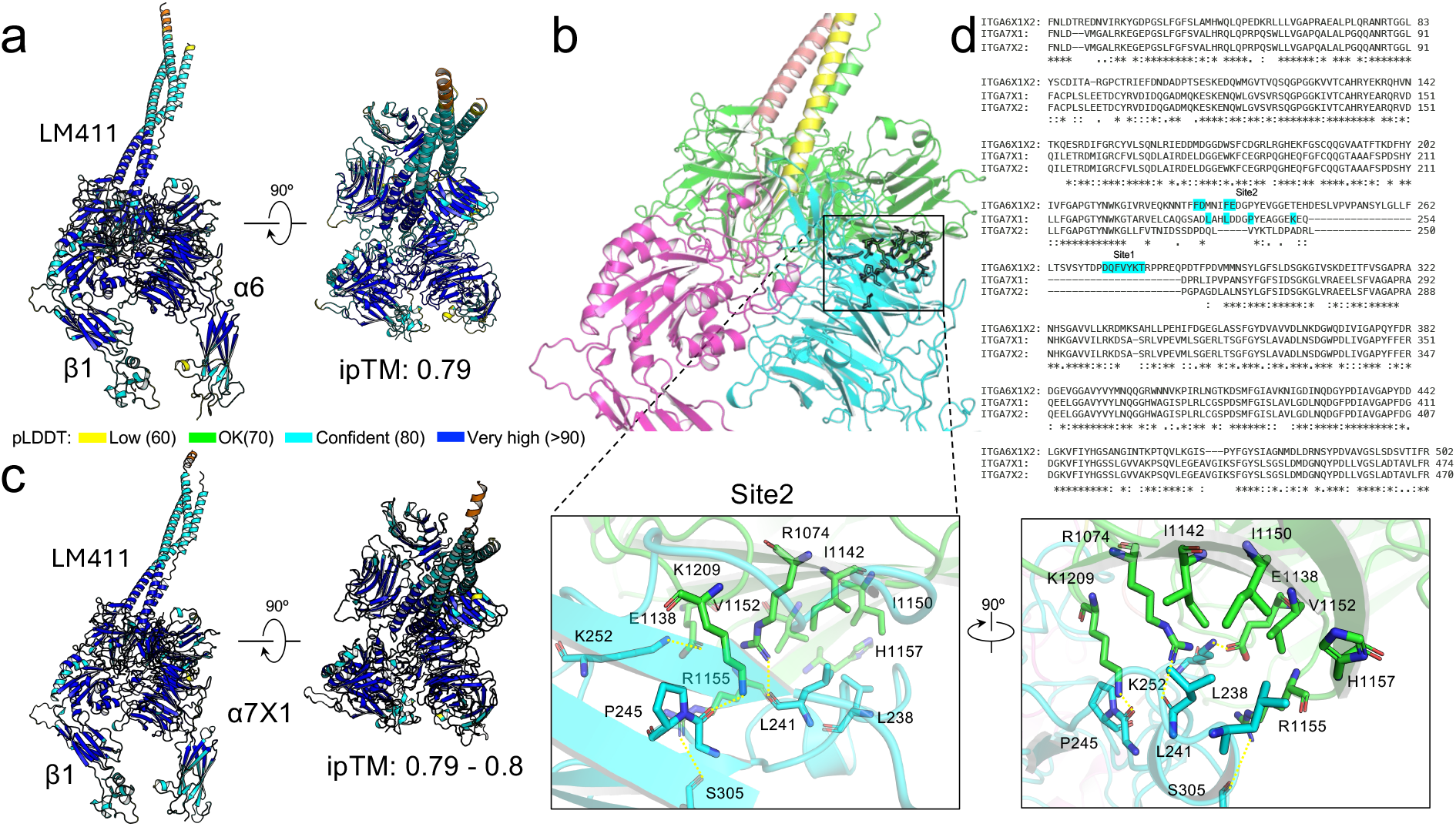
AlphaFold 3 predicted structural models of integrins-T8-LM411 complex. **a**, pLDDT plot of AlphaFold 3 predicted integrin α6β1-T8LM411 complex structure and its ipTM. **b**, AlphaFold 3 predicted structural model of T8-LM411 and integrin α7×1β1 complex and the enlarged detailed view of site2 interactions between integrin α7×1 and LamA4 LG2 domain. **c**, pLDDT plot of AlphaFold 3 predicted integrin α7×1β1-T8LM411 complex structure and its ipTM. **d**, sequence alignment among integrin α6×1×2, α7×1 and α7×2. Residues highlighted in cyan are predicted to interact with LamA4.

**Supplementary Fig 7.**
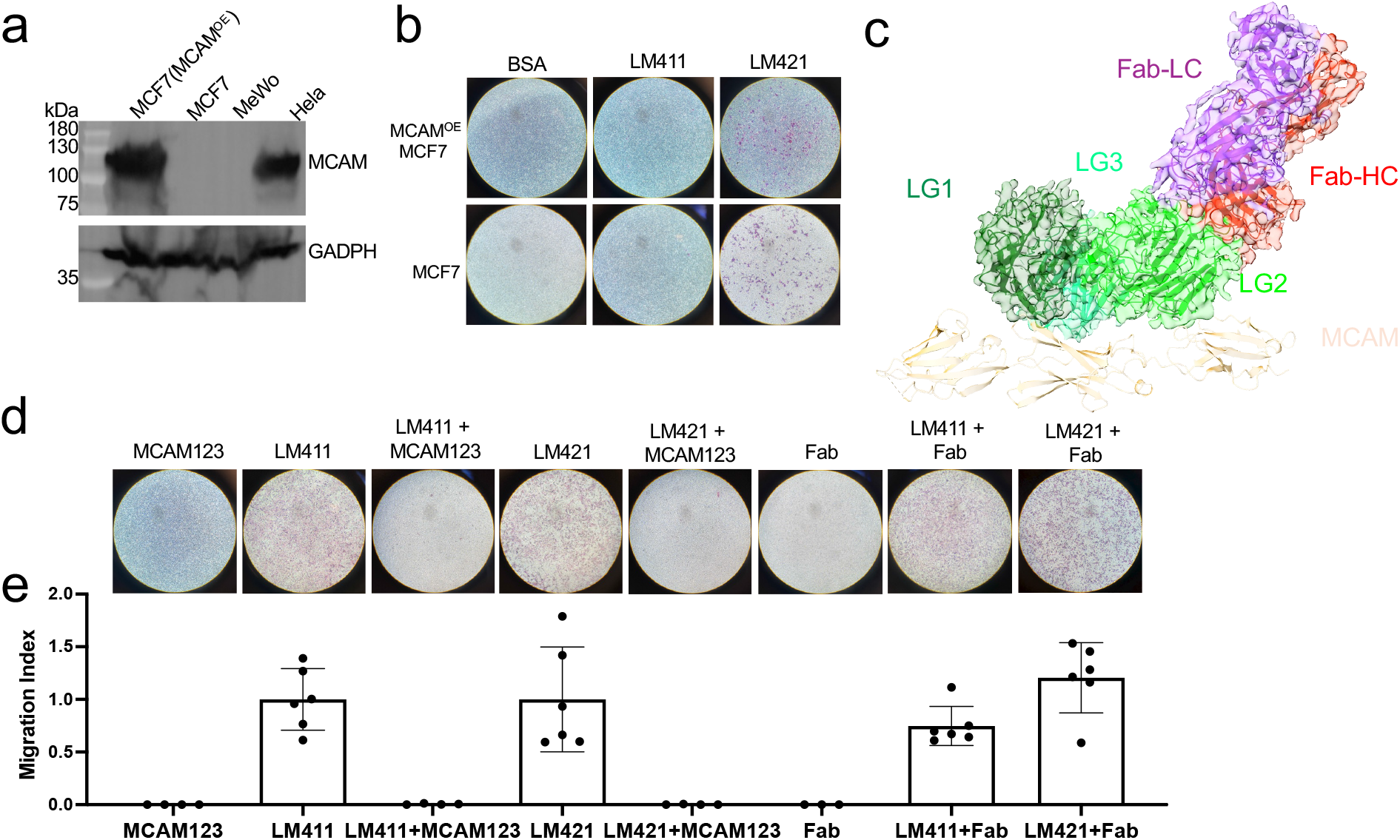
Laminin α4 promotes cell migration primarily through integrin-dependent mechanisms. **a**, Western blot analysis of MCAM protein expression in MCF7 breast cancer cells stably overexpressing full-length MCAM, parental MCF7 cells, and MeWo melanoma cells, with Hela cells included as a positive control. **b**, Overexpression of full-length MCAM in MCF7 breast cancer cells doesn’t endow MCF7 with enhanced migration towards LM411 and LM421. The concentration of LM411 and LM421 is 250 nM. **c**, Surface view of the Cryo-EM reconstruction (3.52 Å, Supplementary Fig. 8) and ribbon diagram of an atomic model of LamA4 LG1-3-Fab complex. MCAM is shown in cartoon representation to illustrate the location of the Fab epitope relative to the MCAM binding interface. **d**, The first three Ig-like domains of MCAM, i.e. MCAM123, which engage in LamA4, are sufficient to inhibit Laminin α4-mediated MeWo melanoma cell migration. In contrast, the Fab recognizing the LG2 domain of LamA4 at an epitope distinct from the MCAM/integrin binding site does not inhibit LamA4-driven MeWo cell migration. The concentrations of LM411 and LM421 are 100 nM and 25 nM, respectively. The concentration of MCAM123 and Fab is 1 μM. e, Quantification of the migration assays shown in d. Data are compiled from at least three independent experiments.

**Supplementary Fig 8.**
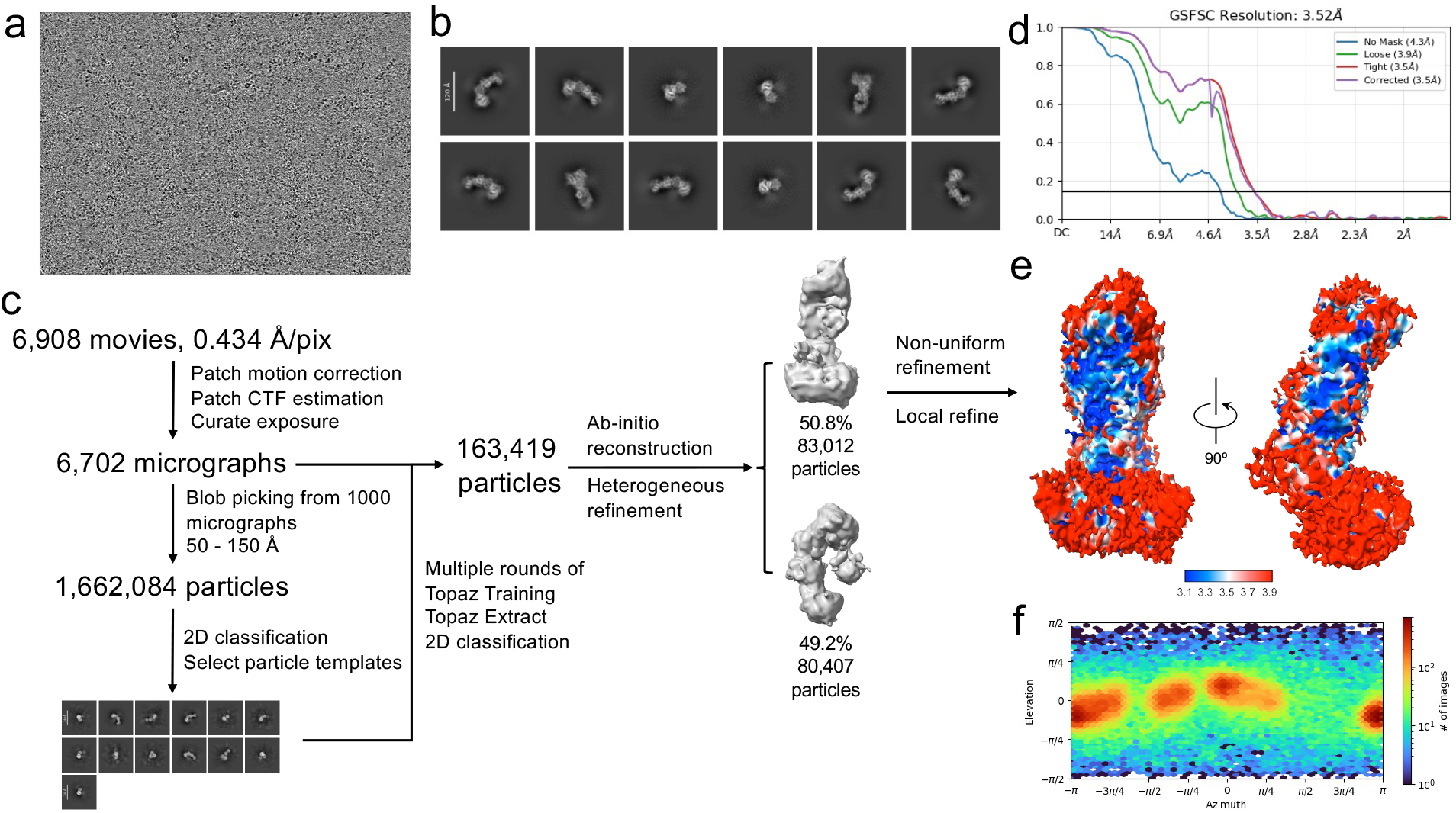
Cryo-EM data processing for the LamA4 LG1-3-Fab complex structure. **a**, Representative raw micrograph of LamA4 LG1-3-Fab complex. **b**, Representative 2D class averages of LamA4 LG1-3-Fab complex. **c**, Images of the processing workflow for structure determination of LamA4 LG1-3-Fab complex. **d**, Gold-standard Fourier Shell Correlation (FSC) of the local-refined map. **e**, Local resolution estimation of the map. **f**, Angular distribution of particles in the final map obtained by non-uniform refinement.

**Supplementary Data Table 1.** Cryo-EM data collection, refinement and validation statistics.

|  | sMCAM –<br>T8-LM411 | sMCAM –<br>LamA4 LG1-3 | sMCAM –<br>T8-LM421 |
| --- | --- | --- | --- |
| **Data collection** |  |  |  |
| Microscope | Glacios | Glacios | Glacios |
| Voltage (kV) | 200 | 200 | 200 |
| Magnification | 45,000 | 45,000 | 45,000 |
| Detector | K3 | K3 | K3 |
| Electron exposure (e<sup>-</sup>/ Å<sup>2</sup>) | 40.87 | 40.2 | 40.95 |
| Defocus range (μm) | -0.8 to -2.0 | -0.8 to -2.0 | -0.8 to -2.0 |
| Pixel size (Å/pixel) | 0.434 | 0.434 | 0.434 |
| **Cryo-EM reconstruction** |  |  |  |
| Particles refined | 184,328 | 319,692 | 81,998 |
| Resolution (Å) | 4.13 | 3.22 | 3.76 |
| FSC threshold | 0.143 | 0.143 | 0.143 |
| Symmetry imposed | C1 | C1 | C1 |
| **Refinement** |  |  |  |
| No. of non-hydrogen atoms | 7778 | 6739 | 7932 |
| Protein residues | 978 | 847 | 998 |
| MolProbity score | 2.32 | 2.16 | 2.03 |
| Clashscore | 20.23 | 18.57 | 13.06 |
| R.m.s. deviations |  |  |  |
| Bond lengths (Å) | 0.003 | 0.004 | 0.003 |
| Bond angles (°) | 0.713 | 0.710 | 0.651 |
| Ramachandran analysis |  |  |  |
| Favored (%) | 90.89 | 94.10 | 93.93 |
| Allowed (%) | 9.01 | 5.78 | 6.07 |
| Outlier (%) | 0.1 | 0.12 | 0.00 |
| Rotamer outlier (%) | 0.00 | 0.4 | 0.34 |

## References

1. Valastyan, S. and R.A. Weinberg, Tumor metastasis: molecular insights and evolving paradigms. Cell, 2011. 147(2): p. 275–92.

2. Lambert, A.W., D.R. Pattabiraman, and R.A. Weinberg, Emerging Biological Principles of Metastasis. Cell, 2017. 168(4): p. 670–691.

3. Reymond, N., B. B. d’Agua, and A.J. Ridley, Crossing the endothelial barrier during metastasis. Nat Rev Cancer, 2013. 13(12): p. 858–70.

4. Yurchenco, P.D., Basement membranes: cell scaffoldings and signaling platforms. Cold Spring Harb Perspect Biol, 2011. 3(2).

5. Hohenester, E. and P.D. Yurchenco, Laminins in basement membrane assembly. Cell Adh Migr, 2013. 7(1): p. 56–63.

6. Sorokin, L., The impact of the extracellular matrix on inflammation. Nat Rev Immunol, 2010. 10(10): p. 712–23.

7. Wu, C., et al., Endothelial basement membrane laminin alpha5 selectively inhibits T lymphocyte extravasation into the brain. Nat Med, 2009. 15(5): p. 519–27.

8. Song, J., et al., Endothelial Basement Membrane Laminin 511 Contributes to Endothelial Junctional Tightness and Thereby Inhibits Leukocyte Transmigration. Cell Rep, 2017. 18(5): p. 1256–1269.

9. Nishiuchi, R., et al., Ligand-binding specificities of laminin-binding integrins: a comprehensive survey of laminin-integrin interactions using recombinant alpha3beta1, alpha6beta1, alpha7beta1 and alpha6beta4 integrins. Matrix Biol, 2006. 25(3): p. 189–97.

10. Arimori, T., et al., Structural mechanism of laminin recognition by integrin.Nat Commun, 2021. 12(1): p. 4012.

11. Lehmann, J.M., G. Riethmuller, and J.P. Johnson, MUC18, a marker of tumor progression in human melanoma, shows sequence similarity to the neural cell adhesion molecules of the immunoglobulin superfamily. Proc Natl Acad Sci U S A, 1989. 86(24): p. 9891–5.

12. Sers, C., G. Riethmuller, and J.P. Johnson, MUC18, a melanoma-progression associated molecule, and its potential role in tumor vascularization and hematogenous spread. Cancer Res, 1994. 54(21): p. 5689–94.

13. Johnson, J.P., et al., Melanoma progression-associated glycoprotein MUC18/MCAM mediates homotypic cell adhesion through interaction with a heterophilic ligand. Int J Cancer, 1997. 73(5): p. 769–74.

14. Wang, Z., et al., CD146, from a melanoma cell adhesion molecule to a signaling receptor. Signal Transduct Target Ther, 2020. 5(1): p. 148.

15. Flanagan, K., et al., Laminin-411 is a vascular ligand for MCAM and facilitates TH17 cell entry into the CNS. PLoS One, 2012. 7(7): p. e40443.

16. Bardin, N., et al., CD146 and its soluble form regulate monocyte transendothelial migration. Arterioscler Thromb Vasc Biol, 2009. 29(5): p. 746–53.

17. Stalin, J., et al., Soluble melanoma cell adhesion molecule (sMCAM/sCD146) promotes angiogenic effects on endothelial progenitor cells through angiomotin. J Biol Chem, 2013. 288(13): p. 8991–9000.

18. Mannion, A.J., et al., Pro- and anti-tumour activities of CD146/MCAM in breast cancer result from its heterogeneous expression and association with epithelial to mesenchymal transition. Front Cell Dev Biol, 2023. 11: p. 1129015.

19. Ishikawa, T., et al., Laminins 411 and 421 differentially promote tumor cell migration via alpha6beta1 integrin and MCAM (CD146). Matrix Biol, 2014. 38: p. 69–83.

20. Ishikawa, T., et al., Monoclonal antibodies to human laminin alpha4 chain globular domain inhibit tumor cell adhesion and migration on laminins 411 and 421, and binding of alpha6beta1 integrin and MCAM to alpha4-laminins. Matrix Biol, 2014. 36: p. 5–14.

21. Aumailley, M., et al., A simplified laminin nomenclature. Matrix Biol, 2005. 24(5): p. 326–32.

22. Takizawa, M., et al., Mechanistic basis for the recognition of laminin-511 by alpha6beta1 integrin. Sci Adv, 2017. 3(9): p. e1701497.

23. Punjani, A., H. Zhang, and D.J. Fleet, Non-uniform refinement: adaptive regularization improves single-particle cryo-EM reconstruction. Nat Methods, 2020. 17(12): p. 1214–1221.

24. Punjani, A., et al., cryoSPARC: algorithms for rapid unsupervised cryo-EM structure determination. Nature Methods, 2017. 14(3): p. 290–+.

25. Abramson, J., et al., Accurate structure prediction of biomolecular interactions with AlphaFold 3. Nature, 2024. 630(8016): p. 493–500.

26. Yamada, M. and K. Sekiguchi, Molecular Basis of Laminin-Integrin Interactions. Curr Top Membr, 2015. 76: p. 197–229.

27. von der Mark, H., et al., Alternative Splice Variants of α7β1 Integrin Selectively Recognize Different Laminin Isoforms. Journal of Biological Chemistry, 2002. 277(8): p. 6012–6016.

28. Goodman, S.L. and M. Picard, Integrins as therapeutic targets. Trends Pharmacol Sci, 2012. 33(7): p. 405–12.

29. Raab-Westphal, S., J.F. Marshall, and S.L. Goodman, Integrins as Therapeutic Targets: Successes and Cancers. Cancers (Basel), 2017. 9(9).

30. Zondler, L., et al., MCAM/CD146 Signaling via PLCγ1 Leads to Activation of β1-Integrins in Memory T-Cells Resulting in Increased Brain Infiltration. Frontiers in Immunology, 2020. 11.

31. Bepler, T., et al., Positive-unlabeled convolutional neural networks for particle picking in cryo-electron micrographs. Nature Methods, 2019. 16(11): p. 1153–+.

32. Liebschner, D., et al., Macromolecular structure determination using X-rays, neutrons and electrons: recent developments in Phenix. Acta Crystallogr D Struct Biol, 2019. 75(Pt 10): p. 861–877.

33. Afonine, P.V., et al., Real-space refinement in PHENIX for cryo-EM and crystallography. Acta Crystallogr D Struct Biol, 2018. 74(Pt 6): p. 531–544.

34. Emsley, P., et al., Features and development of Coot. Acta Crystallogr D Biol Crystallogr, 2010. 66(Pt 4): p. 486–501.

